# Working memory processing boosts the neural representation of long-term memories

**DOI:** 10.1101/2024.08.15.608072

**Authors:** Melinda Sabo, Laura-Isabelle Klatt, Daniel Schneider

## Abstract

Understanding the interaction between short-term and long-term memory systems is essential for advancing our knowledge of human memory. This study investigates whether working memory processes, specifically attentional prioritization (Experiments 1 and 2) and testing (Experiment 2), can enhance the neural activity associated with recently encoded long-term memories. A total of 86 participants completed a novel three-phase memory task that integrated a traditional long-term memory learning task with a working memory paradigm. In the first phase, participants encoded object-location associations. During the second phase, these associations were reintroduced in a working memory task that manipulated attentional prioritization; participants were required to report the location of the cued object. In the final phase, participants recalled the locations associated with each object. By analyzing both behavioral performance and electroencephalogram (EEG) data collected during this retrieval phase, we found that attentional prioritization in working memory significantly improved long-term memory retrieval, a finding supported by corresponding neural evidence. Additionally, Experiment 2 demonstrated that both prioritization and testing in working memory jointly enhance the neural representation of long-term memories. These findings indicate that working memory processes can dynamically alter the neural patterns underlying long-term memory representations, revealing a more integrated role for working memory in long-term memory consolidation.

## 1. Introduction

Understanding the dynamic interplay between our short-term and long-term memory systems is essential in the pursuit of understanding human memory and learning. Already in the 1960s, theories proposed by Atkinson and Shiffrin suggested a continuous interaction between these memory systems^1^. These ideas remain influential in contemporary memory research. For instance, Cowan and colleagues propose a simultaneous information exchange between working-and long-term memory. Accordingly, information in working memory activates long-term memory representations^2,3^, while retrieved long-term memory content is temporarily maintained in working memory^4^. Likewise, Oberauer and colleagues discuss the existence of a flexible gate mechanism that regulates the entry of long-term memory content into working memory, allowing it only when beneficial^5–8^. This reciprocal exchange underscores the active role of working memory in shaping and accessing long-term memory representations.

Previous research has extensively explored how various maintenance and manipulation processes in working memory affect the transfer of information to long-term memory and its subsequent recall. A key phenomenon in this area is the McCabe effect, which shows improved long-term recall for words interrupted by an arithmetic task during working memory maintenance compared to uninterrupted words (complex vs. simple span tasks)^9^. This improvement is attributed to the covert retrieval of words during the maintenance period while solving the arithmetic task^9–11^ (for conflicting evidence obtained in experiments using complex stimuli, see^12^). Other explanations include benefits from the extended time spent in working memory^13,14^ or elaboration during the maintenance period^15^.

Studies on the transfer of information between working memory and long-term memory have also examined the role of attentional processes. Depending on their operationalization, researchers have assessed how attentional refreshing, value-based attentional prioritization, and cue-based attentional prioritization in working memory affect information transfer to long-term memory and subsequent retrieval. Attentional refreshing, which involves briefly thinking about a recently encountered stimulus to reactivate its representation^16^, shows mixed evidence regarding its benefit for delayed long-term memory recall. Some studies suggest that attentional refreshing in working memory leads to enhanced long-term memory retrieval^17,18^, while others do not find such benefits^19^ beyond simple repetition^20^. In contrast, the effects of attentional prioritization on long-term memory retrieval depend on the type of cue. Cue-based attentional prioritization in working memory has been linked to enhanced long-term recall^21–25^, whereas reward-cue-based prioritization was not associated with such long-term benefits^25,26^. These findings suggest that the operationalization of attentional processes matters when studying long-term memory recall benefits, with cue-based attentional prioritization proving to be the most robust effect.

The studies described above consistently examined how attentional processes in working memory impact the transfer of information from short-term to long-term memory. However, an important yet unanswered question is whether these working memory processes can also enhance *existing* long-term memory representations that have been recently encoded but not yet consolidated. This question is crucial because evidence of such an effect would suggest that working memory not only facilitates the transfer of information to long-term memory but also actively modifies and strengthens existing representations. This implies a more dynamic and flexible relationship between working memory and long-term memory than previously thought. Instead of merely acting as a temporary buffer, working memory could also serve as an instance, in which long-term memories are further manipulated and reinforced. These findings would significantly expand our understanding of the interaction between these two memory systems.

In Experiment 1, we examined the extent to which cue-based attentional prioritization in working memory enhances the retrieval of previously encoded long-term memory representations. We investigate this effect not only behaviorally, but also at the neural level via the parietal old-new effect, an established EEG component of long-term memory recollection^27–31^. To foreshadow the results of Experiment 1, we found that both behavioral and EEG data supported enhanced long-term memory retrieval of information undergoing cued- based attentional prioritization in working memory.

To determine whether the observed effect is due solely to attentional prioritization or a combination of attentional prioritization and testing effects in working memory, we conducted Experiment 2. Both behavioral and neural evidence revealed that, compared to a control condition, items that were either prioritized or tested in working memory showed enhanced long-term memory retrieval. Similarly, EEG results suggested that both attentional prioritization and testing in working memory enhance long-term memory representations and their retrieval. Importantly, accuracy data revealed a distinct benefit for attentional prioritization over testing, a difference not reflected at the neural level. Overall, findings are discussed in light of attentional prioritization and testing in working memory *jointly* contributing to the enhancement of long-term memory representations and their subsequent retrieval. Finally, we discuss the theoretical implications of these results for existing theories describing the relationship between long-term- and working memory.

## 2. Results

In Experiment 1 participants completed a combined episodic long-term and working memory task with three phases (see Figure 1a and b). In the first phase, they learned object-location associations with the instruction to remember them for later recall. In the second phase, a working memory (retroactive cuing / retro-cue) paradigm was introduced. Participants were shown objects from the first phase, presented on their learned locations, along with two task- irrelevant scrambled objects. Participants’ task was to memorize the objects for a subsequent test. After a short maintenance period, a 100% informative retro-cue indicated which object would be tested. During test, participants had to decide if the cued object matched a centrally presented object (i.e., probe). Each object appeared four times on the same location during the working memory task. In the final retrieval phase, participants were shown all objects from the first phase and were asked to report the location learnt in the first phase using a response device with four spatially aligned buttons corresponding to the four possible locations (see Figure 1b). All analyses were conducted on data from this third, long-term memory retrieval phase. Notably, for our analysis these trials were sorted based on the processing that occurred during the working memory task (i.e., the second phase), which determined the experimental condition (for an overview, of the conditions, see Figure 1c).

**Figure 1.**
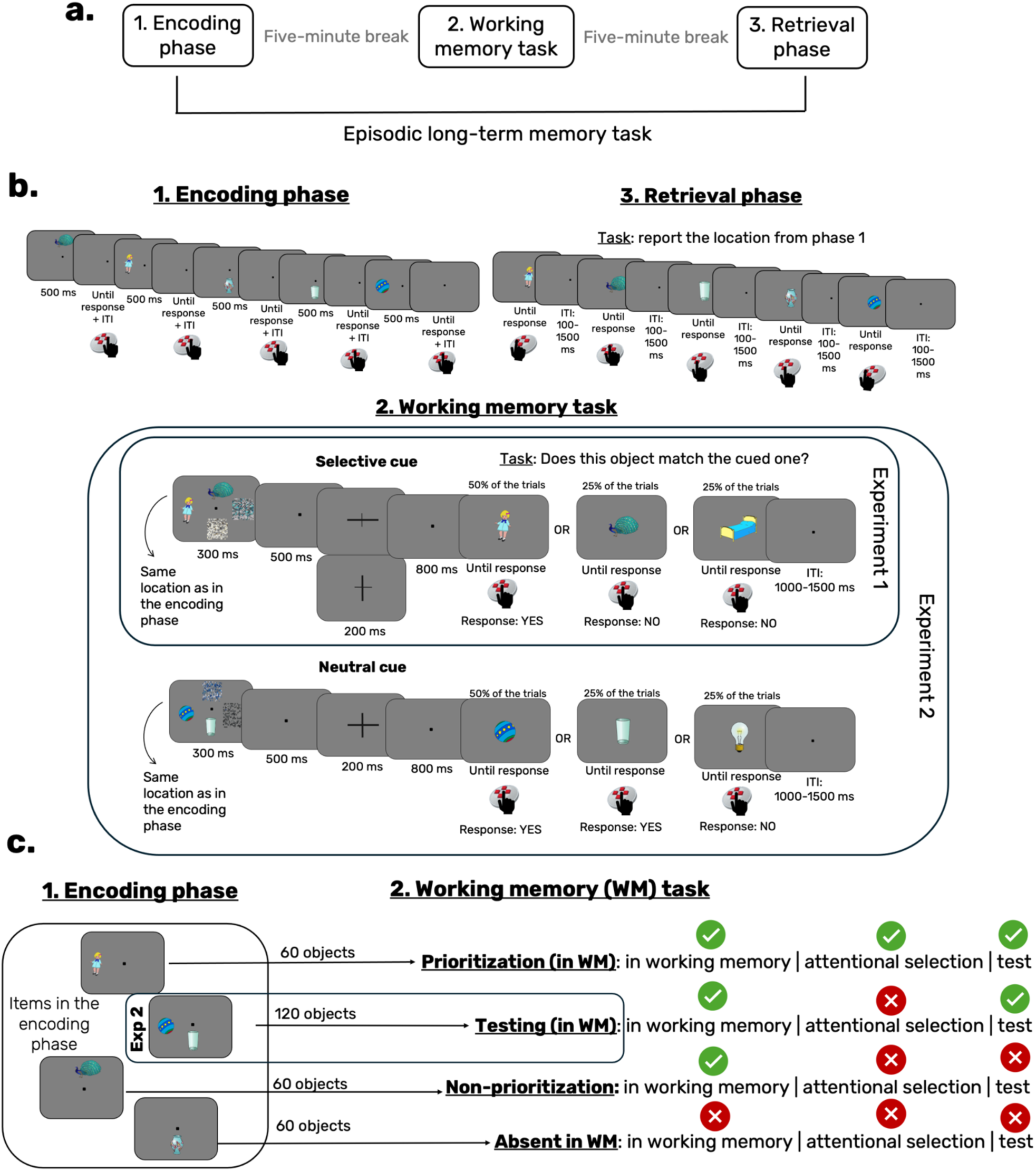
Overview of the experimental design. Panel a outlines the three main phases of the experiment, while Panel b offers a detailed breakdown of these phases. In Experiment 1, the working memory phase includes only selective (informative) retro-cues, whereas Experiment 2 incorporates both selective cues and neutral (non-informative) cues. Panel c demonstrates how the same objects from the encoding phase reappear during the working memory task, undergoing different types of processing depending on their assigned condition. For instance, in the “prioritization in WM” condition, the object undergoes both attentional selection and testing within working memory, while in the “absent in WM” condition, the object is not presented during the working memory task, appearing only during encoding and the final retrieval phase. Notably, the “testing in WM” condition is exclusive to Experiment 2, whereas the other three conditions are common to both experiments.

Experiment 1 includes three experimental conditions based on how objects were processed in working memory (or if they were excluded from such processing): (i) the *prioritization condition*, involves attentional prioritization and testing of the previously encoded long-term memory representation in working memory; (ii) the *non-prioritization condition*, includes a brief presentation and maintenance of the object followed by the item being placed outside the focus of attention, as it is no longer relevant for the subsequent working memory test (also analogous to a repetition effect); and (iii) *absent in working memory* (control) *condition*, in which the object is initially encoded into long-term memory and retrieved in the last phase, but does not appear in the working memory task.

According to our hypothesis, if attentional prioritization in working memory enhances *existing* long-term memory representations that have been recently encoded but not yet consolidated, we should observe improved behavioral performance and a stronger parietal old-new effect^27–31^ in the prioritization compared to the non-prioritization and absent in working memory condition. This would suggest that cue-based attentional prioritization in working memory has the capacity to enhance the recall of long-term memories and that this effect goes beyond simple repetition within working memory (non-prioritization condition).

### 2.1. Long-term memory accuracy is boosted by attentional prioritization in working memory – evidence from Experiment 1

Table 1 shows the average retrieval phase accuracy, indicating that participants were most accurate in the prioritization condition, followed by the non-prioritization condition, and least accurate in the absent in working memory condition. Statistical comparisons using repeated measures ANOVA (rm-ANOVA) revealed that sphericity was violated (*χ*^2^(42) = 7.58, *p* = .02, ε = 0.85), so Greenhouse–Geisser corrected results are reported. The accuracy analysis showed a main effect of condition: *F* (2, 84) = 51.85, *pcorr* < .001, 11p^2^= 0.55, *BF10* = 1.98×10^12, with the Bayes factor indicating very strong evidence in favor of the alternative hypothesis (see Figure 2a). The Bayes factor of the post-hoc analyses revealed very strong evidence for the accuracy difference between: (i) the prioritization vs. non-prioritization condition: *t*(42) = 6.51, *padj* < .001, *dav* = 0.70, 95% CI [7.29, 13.83], *BF10 corr* = 116301.91; (ii) the prioritization vs. absent in working memory condition: *t*(42) = 8.68, *padj* < .001, *dav* = 1.05, 95% CI [12.10, 19.44], *BF10 corr* = 9.13×10^7; (iii) the non-prioritization vs. absent in working memory condition: *t*(42) = 4.19, *padj* < .001, *dav* = 0.29, 95% CI [2.70, 7.72], *BF10 corr* = 104.60. These results indicate that cued attentional prioritization in working memory led to an increase in long-term memory accuracy, surpassing performance for both items that were absent in the working memory task and those that were merely presented (but never tested) in the working memory task.

**Figure 2.**
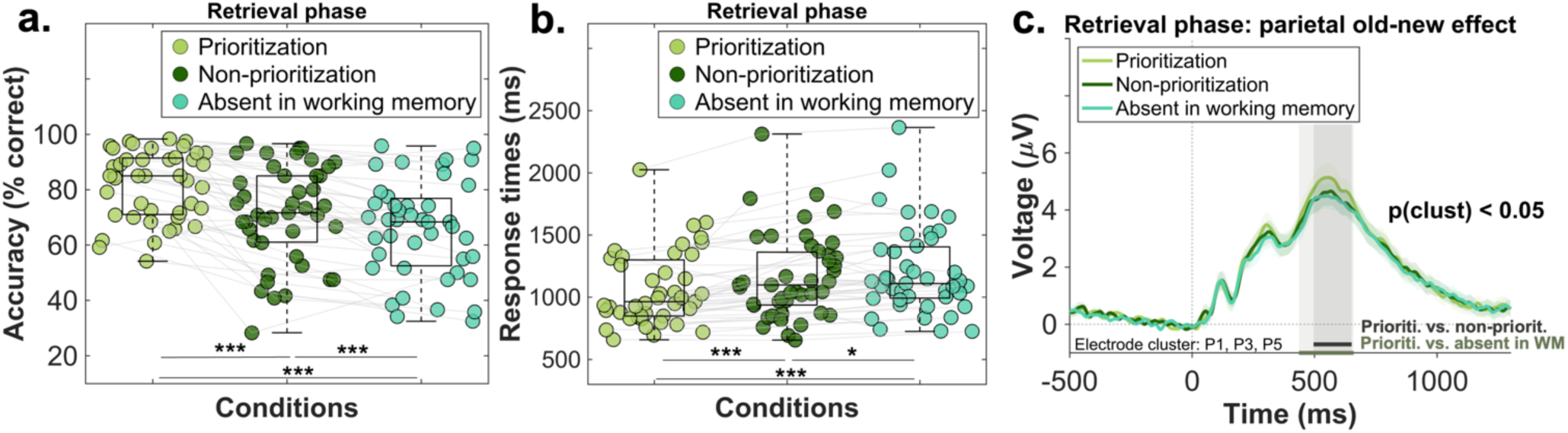
Results of Experiment 1. The scatterplots of panel a and b illustrate the mean accuracy and mean reaction time recorded during the three conditions of the final long-term memory retrieval phase. The central mark in each boxplot represents the median, while the bottom and top edges denote the 25th and 75th percentiles, respectively. The whiskers do not extend to averages, which are treated as outliers. Grey lines connect data points from the same participant. Statistical significance is indicated with the following symbols: \**p* < 0.05; \*\**p* < 0.01, \*\*\**p* < 0.001. Panel c illustrates the parietal old-new effect recorded during the same retrieval phase. The three shaded regions indicate the significant time window identified through the cluster-based permutation analysis: (i) 496-652 ms for the prioritization vs. non- prioritization contrast and (ii) 436-660 ms for the prioritization vs. absent in working memory condition contrast. The shaded area around the ERP of each condition denotes the standard error of the mean.

**Table 1.** Descriptive statistics – Experiment 1.

and standard deviation (SD)
| **Condition** | **N** | **Mean accuracy** | **SD accuracy** | **Mean response times** | **SD response times** |
| --- | --- | --- | --- | --- | --- |
| prioritization | 43 | 81.68% | 12.39% | 1060.80 ms | 294.63 ms |
| non-prioritization | 43 | 71.12% | 17.60% | 1166.81 ms | 342.96 ms |
| absent in working memory | 43 | 65.91% | 17.58% | 1206.22 ms | 350.33 ms |

### 2.2. Decreased response times during long-term memory retrieval for prioritized items in working memory – evidence from Experiment 1

Descriptive analysis (see Table 1) showed that response times in the long-term memory retrieval phase followed the same pattern as accuracy: participants were fastest for items that appeared in the prioritization condition during the working memory task, followed by the non-prioritization condition, and slowest in the absent in working memory condition. A comparable repeated measures ANOVA (rm-ANOVA) was conducted to compare response times across conditions. Since sphericity was violated (*χ^2^*(42) = 14.03, *p* < .001, ε = 0.77), Greenhouse– Geisser corrected results are reported. Results of the rm-ANOVA indicated a main effect of condition: *F* (2, 84) = 27.32, *pcorr* < .001, 11p^2^= 0.39, *BF10* = 1.17×10^7, with the Bayes factor indicating very strong evidence in favor of the alternative hypothesis (see Figure 2b). The Bayes factor of the post-hoc analyses suggested very strong evidence for the response times difference between: (i) the prioritization vs. non-prioritization condition: *t*(42) = -5.23, *padj* < .001, *dav* = 0.33, 95% CI [-146.89, -65.12], *BF10 corr* = 2213.76; and (ii) the prioritization vs. absent in working memory condition: *t*(42) = -5.89, *padj* < .001, *dav* = 0.45, 95% CI [-195.21,95.62], *BF10 corr* = 16866.52. The response times difference between the non-prioritization vs. absent in working memory condition was only supported by weak evidence: *t*(42) = -2.64, *padj* = .01, *dav* = 0.11, 95% CI [-69.53, -9.29], *BF01 corr* = 2.06. Overall, similar to the accuracy results, the response times pattern also confirms that cued prioritization is associated with faster responses, which exceed the benefits observed when the item is either absent from working memory or briefly presented and maintained during the working memory task.

### 2.3. Enhanced parietal old-new component for prioritized items in working memory – evidence from Experiment 1

In addition to the behavioral measures, we investigated whether modulations of the parietal old-new effect mirrored the observed behavioral patterns. This event-related potential component is typically measured in tasks requiring old (previously encountered) versus new (never encountered) decisions during memory retrieval and has been argued to reflect the conscious recollection of long-term memories^28,30–32^. Previous research has also shown that this component can be differentially modulated by the retrieval of deeply versus superficially encoded information^29^ or by the retrieval of information with varying precision levels^33^. As indicated in Figure 2c, data from the electrode cluster P1, P3, P5^34^ was averaged and contrasted between conditions via cluster-based permutation statistics. Our analysis revealed a significant cluster between 496-652 ms for the prioritization vs. non-prioritization contrast and a cluster between 436-660 ms for the prioritization vs. absent in working memory condition contrast. No statistically reliable effects were found when the non-prioritization and absent in working memory condition were compared. These results indicate an enhanced long-term memory retrieval for the object-location associations that underwent attentional selection in working memory compared to the other two conditions.

### 2.4. Overview of Experiment 2

As outlined in the Introduction, Experiment 2 aimed to determine whether the effects observed in Experiment 1 were due solely to attentional prioritization or also to testing effects in working memory. We define the testing effect in this context as the extended, non-selective maintenance of information in working memory to generate a response during the final match-no-match test. However, for brevity, we will use the term “testing or testing effect”. To disentangle attentional prioritization from testing effects, we introduced a new type of cue in the working memory task: uninformative/neutral cues. Similar to Experiment 1, participants encoded the stimulus display and were probed in a match-no-match task at the end of the trial. However, unlike informative cues, neutral cues did not provide information about the to-be-tested item, requiring participants to maintain both objects for the comparison in the working memory test. Importantly, similar to Experiment 1, participants did not report any locations during the working memory task. All other aspects were kept consistent with Experiment 1.

As illustrated in Figure 1c, Experiment 2 assumes that objects in the *prioritization condition* undergo both attentional prioritization and testing in working memory. In contrast, objects in the *testing condition* undergo a non-selective maintenance and testing at the end of the trial. By comparing the final long-term memory retrieval of objects between these two conditions, we can determine whether the benefits observed in Experiment 1 are due solely to attentional prioritization or a combination of attentional prioritization and testing. The other two conditions (*non-prioritization in working memory* and *absent in working memory*) are identical to Experiment 1. According to our hypothesis, if long-term memory performance and the amplitude of the parietal old-new component are higher in the prioritization condition compared to the testing condition (with both differing from the absent in working memory condition and non-prioritization condition), it would suggest that attentional prioritization provides an additional long-term memory benefit beyond testing.

### 2.5. Long-term memory accuracy is boosted by attentional prioritization above and beyond testing in working memory – evidence from Experiment 2

At a descriptive level (see Table 2), participants were the most accurate in the prioritization condition, followed by the testing condition, then in the non-prioritization condition, and finally in the absent in working memory condition. To contrast participant’s accuracy across the experimental conditions, we conducted a rm-ANOVA. Because the sphericity assumption was violated (*χ^2^*(42) = 21.42, *p* < .001, ε = 0.75), the Greenhouse–Geisser correction was applied. Results suggested a main effect of condition, which was also supported by the very strong evidence in favor of the alternative model as revealed by the Bayes factor: *F* (3, 126) = 63.76, *pcorr* < .001, 11p^2^= 0.60, *BF10* = 6.36×10^21 (Figure 3a). The Bayes factor of the post-hoc tests revealed very strong evidence for the accuracy difference between (i) the prioritization vs. non-prioritization condition: *t*(42) = 11.26, *padj* < .001, *dav* = 0.75, 95% CI [9.76, 14.03], *BF10 corr* = 1.057×10^11; (ii) between the prioritization vs. absent in working memory condition: *t*(42) = 10.62, *padj* < .001, *dav* = 0.93, 95% CI [12.18, 17.89], *BF10 corr* = 1.806×10^10; (iii) between the prioritization vs. testing condition: *t*(42) = 10.52, *padj* < .001, *dav* = 0.52, 95% CI [6.61, 9.74], *BF10 corr* = 1.374×10^10; (iv) between the absent in working memory condition vs. testing condition: *t*(42) = -5.16, *padj* < .001, *dav* = 0.40, 95% CI [-9.54, -4.18], *BF10 corr* = 1280.32.

**Figure 3.**
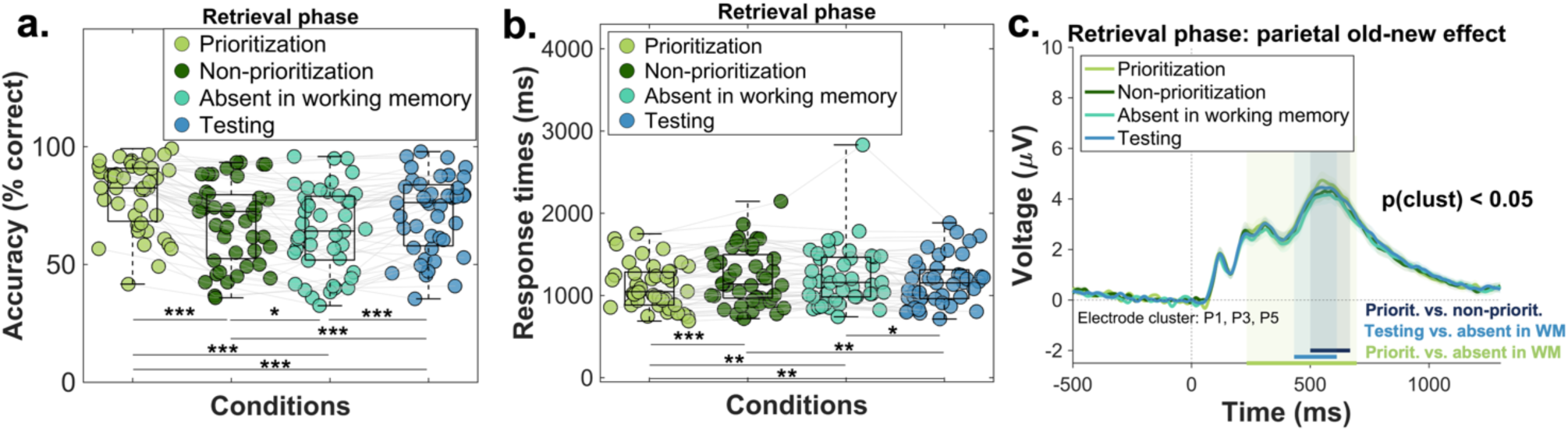
**Behavioral and ERP results of Experiment 2**. The scatterplots of panel a and b illustrate the mean accuracy and mean reaction time recorded during the four conditions of the final long-term memory retrieval phase. The central mark in each boxplot represents the median, while the bottom and top edges denote the 25th and 75th percentiles, respectively. The whiskers do not extend to averages, which are treated as outliers. Grey lines connect data points from the same participant. Statistical significance is indicated with the following symbols: \**p* < 0.05; \*\**p* < 0.01, \*\*\**p* < 0.001. Panel c illustrates the parietal old-new effect recorded during the same retrieval phase. The three shaded regions indicate the significant time window identified through the cluster-based permutation analysis: (i) 500- 668 ms for prioritization vs. non-prioritization, (ii) 232-696 ms for prioritization vs. absent in working memory condition, and (iii) 432-612 ms for testing vs. absent in working memory condition. The shaded areas around the ERP of each condition denote the standard error of the mean.

**Table 2.** Descriptive statistics – Experiment 2.

and standard deviation (SD) – Experiment 2
| **Condition** | **N** | **Mean**<br>**accuracy** | **SD accuracy** | **Mean response**<br>**times** | **SD response times** |
| --- | --- | --- | --- | --- | --- |
| prioritization | 43 | 79.65% | 14.53% | 1108.37 ms | 273.99 ms |
| non-prioritization | 43 | 67.75% | 17.15% | 1220.77 ms | 342.00 ms |
| absent in working<br>memory | 43 | 64.61% | 17.78% | 1229.77 ms | 369.52 ms |
| testing | 43 | 71.47% | 16.46% | 1164.54 ms | 291.11 ms |

**Table 3.** Overview of the outcome of each preprocessing pipeline.

| Rejected number of<br>channels | Number of remaining<br>trials before ICA | Number of rejected<br>IC components | Number of remaining<br>trials after the ICA |
| --- | --- | --- | --- |
| $M = 1.62$ Range: 0-5 | $M = 311.34$ (86.49 %)<br>Range: 253-348<br>(70.27 % - 96.66 %) | $M = 31.97$<br>Range: 14-58 | $M = 312.95$ (86.93 %)<br>Range: 239-358<br>(66.38% - 99.44%) |

**Table 4.**
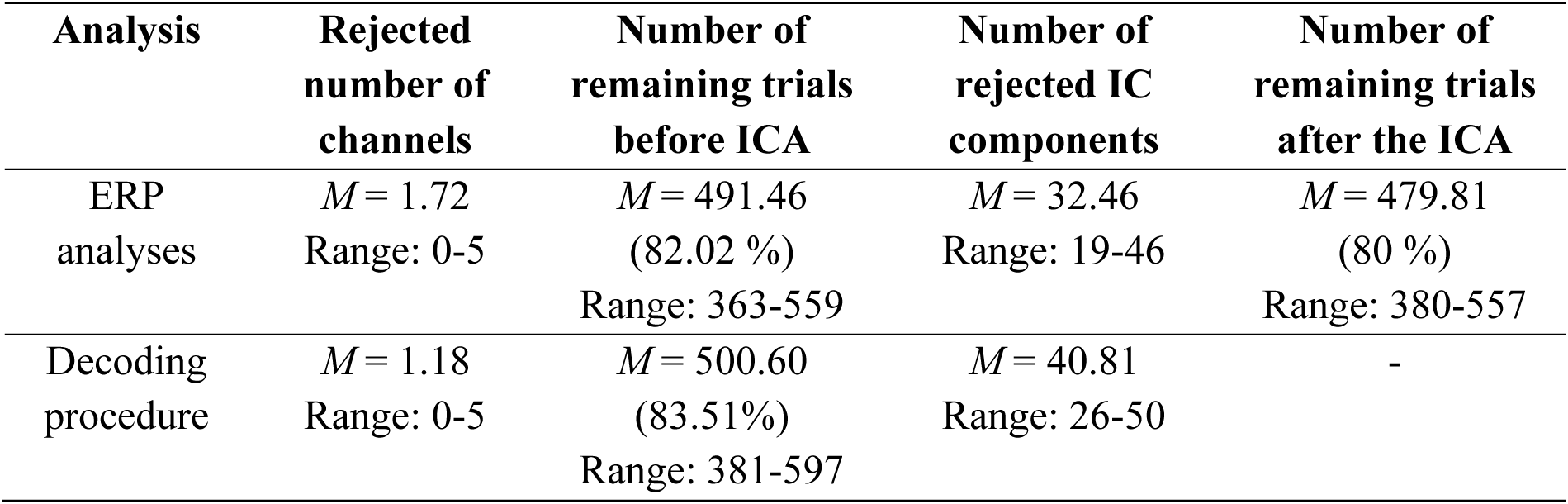
Overview of the outcome of each preprocessing pipeline.

Additionally, strong evidence was found for the comparison between non-prioritization and testing condition *t*(42) = -4.18, *padj* < .001, *dav* = 0.22, 95% CI [-5.51, -1.92], *BF10 corr* = 70.91. Finally, weak evidence was found for the contrast of non-prioritization vs. absent in working memory condition: *t*(42) = 2.44, *padj* = 0.01, *dav* = 0.17, 95% CI [0.54, 5.73], *BF01 corr* = 0.96. Regarding our main hypothesis, the results suggest that both the prioritization and testing condition resulted in higher accuracy compared to the absent in working memory condition and non-prioritization condition. Interestingly, the testing condition still yielded benefits even though participants did not report any locations in this case. Additionally, accuracy was higher in the prioritization condition compared to the testing condition, indicating a greater long-term memory boost for items undergoing attentional prioritization and testing compared to those undergoing only testing in working memory. These findings remain consistent even when the number of times an object appeared as a probe during the match-no-match task is controlled (see Supplementary materials, section 2, Figure S2 for details).

### 2.6. Weak evidence for a response time decrease in the prioritizations vs. testing contrast – evidence from Experiment 2

As indicated in Table 2, at a descriptive level, participants were fastest in the prioritization condition, followed by the testing condition, then in the non-prioritization condition, and finally in the absent in working memory condition. We conducted an rm-ANOVA to investigate the effect of condition on long-term memory response times. Sphericity was again violated (*χ^2^*(42) = 30.13, *p* < .001, ε = 0.65), thus the Greenhouse–Geisser corrected results are reported: *F* (3, 126) = 9.94, *pcorr* < .001, 11p^2^= 0.19, *BF10* = 2773.96. The Bayes factor revealed again very strong evidence for the alternative hypothesis, which predicts a main effect of condition (Figure 3b). The post-hoc tests revealed very strong evidence for the accuracy difference between the prioritization vs. non-prioritization condition: *t*(42) = -4.67, *padj* < .001, *dav* = 0.36, 95% CI [- 160.97, -63.83], *BF10 corr* = 290.72. Similarly, substantial evidence was found supporting the difference between the (i) prioritization vs. absent in working memory condition: *t*(42) = -3.47, *padj* =.003, *dav* = 0.37, 95% CI [-191.97, -50.83], *BF10 corr* = 10.42 and (ii) the non-prioritization vs. testing comparison: *t*(42) = 3.10, *padj* = .006, *dav* = 0.17, 95% CI [19.62, 92.83], *BF10 corr* = 4.14. Weak support was found for the prioritization vs. testing contrast, *t*(42) = -2.94, *padj* = .007, *dav* = 0.19, 95% CI [-94.67, -17.67], *BF10 corr* = 2.86. Finally, our analysis revealed inconclusive evidence for the response times difference between the absent in working memory condition vs. testing condition, *t*(42) = 2.37, *padj* =.02, *dav* = 0.19, 95% CI [9.70, 120.75], *BF10 corr* = 0.83 and substantial evidence for the lack of difference between the non-prioritization vs. absent in working memory condition: *t*(42) = -0.37, *padj* =.70, *dav* = 0.02, 95% CI [-57.21, 39.22], *BF01 corr* = 0.07. Overall, the results indicate that the prioritization condition was associated with faster responses compared to the absent in working memory condition and non- prioritization condition. For the testing condition, faster responses received substantial support only when compared to the non-prioritization condition. The difference between the testing vs. absent in working memory condition was only supported by the frequentist statistical framework. Finally, weak evidence supported faster responses in the prioritization condition compared to the testing condition. Overall, the response time results do not entirely mirror the accuracy results; however, they suggest slightly decreased response times for information undergoing attentional prioritization compared to testing in working memory.

### 2.7. Enhanced parietal old-new component for prioritized and tested items in working memory – evidence from Experiment 2

Similar to Experiment 1, we investigated the parietal old-new effect as an EEG correlate of long-term memory recollection. Data from the electrode cluster P1, P3, P5^34^ were averaged and contrasted across all six condition combinations using cluster-based permutation statistics. The results revealed significant clusters between (i) 500-668 ms for prioritization vs. non-prioritization, (ii) 232-696 ms for prioritization vs. absent in working memory condition, and (iii) 432-612 ms for testing vs. absent in working memory condition (Figure 3c). The other comparisons (prioritization vs. testing, non-prioritization vs. absent in working memory condition, and non-prioritization vs. testing) were not statistically significant. These findings suggest that long-term memory retrieval was enhanced both in the prioritization and in the testing condition when compared to the absent in working memory condition. However, no statistical support was found for the prioritization vs. testing contrast. When it comes to the non-prioritization condition, we could replicate the result pattern from Experiment 1: we found an enhanced parietal old-new effect for prioritization vs. non-prioritization, suggesting that the benefit of attentional prioritization was beyond mere repetition. Additionally, the non-prioritization vs. absent in working memory condition difference was again non-significant. Finally, contrary to our expectation, the non-prioritization vs. testing contrast was not significant, raising the question of whether at the neural level, these two conditions are distinguishable.

### 2.8. Comparable location decoding for prioritized and tested items in working memory – evidence from Experiment 2

Since our ERP analysis showed no differences between the prioritization and testing condition, which contradicts the behavioral results, we aimed to further elucidate the neural mechanisms underlying long-term memory retrieval of locations using a decoding procedure. Previous research suggests that decoding approaches are suitable for tracking how certain information is represented in the brain. Unlike univariate methods, multivariate methods can capture the complex relationship between different input features (e.g., channels, frequencies)^35^. We wanted to investigate whether the location information stored in long-term memory underwent any changes due to differential processing in working memory, with a specific focus on tracking representational changes between the prioritization and testing condition.

As a first step in our analysis, we assessed whether we could decode the location information in each experimental condition. Our classifier (support vector machine algorithm) was trained to distinguish one retrieved location (e.g., left) from all other retrieved locations (i.e., right, top, bottom) during the final retrieval phase. With four possible locations, the chance level is 0.25. As shown in Figure 4a, we found significant above-chance decoding accuracy in the: (i) prioritization condition (significant cluster: 520-836 ms); (ii) testing condition (significant cluster: 88-1100 ms), and (iii) absent in working memory condition (significant cluster: 1028-1200 ms). No statistically reliable effects were found in the non-prioritization condition (Figure 4a). Since we could not successfully decode the location retrieval of the non-prioritization objects, subsequent comparisons were conducted only for the three other conditions.

**Figure 4.**
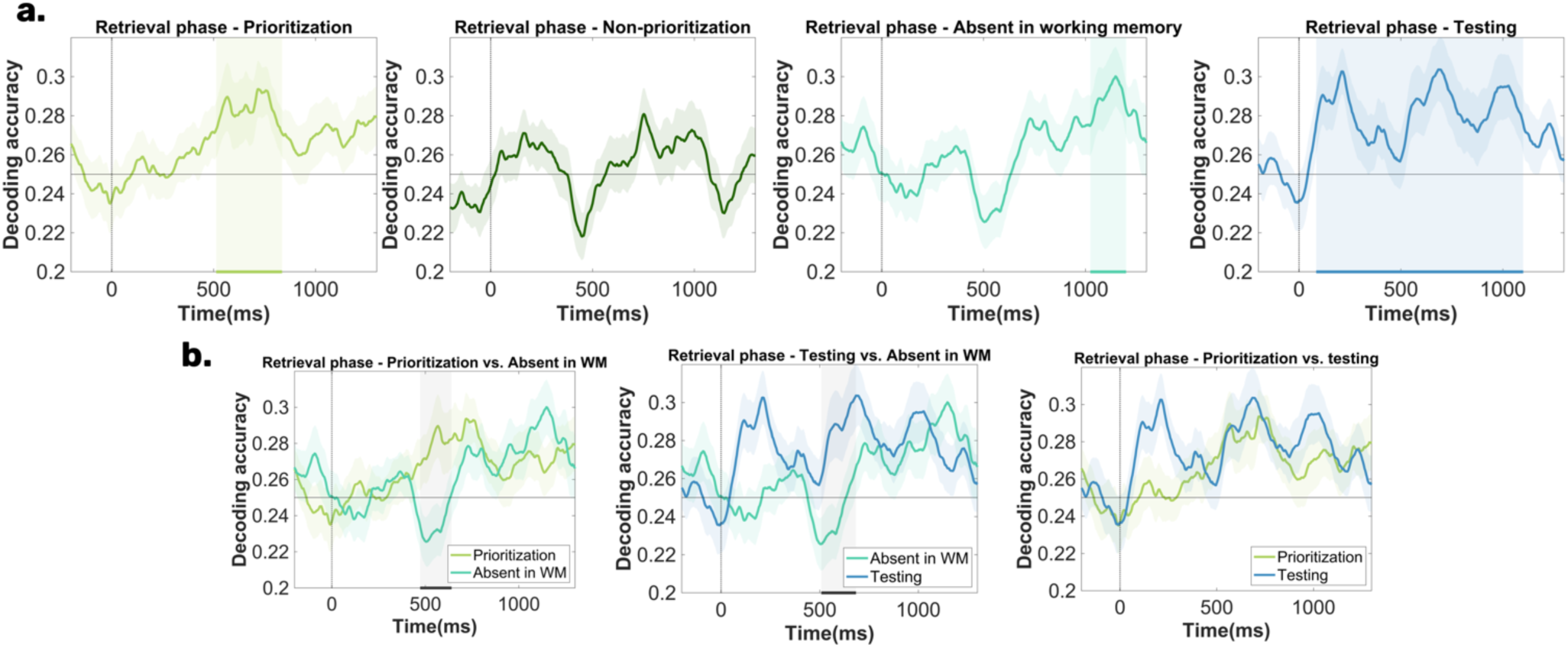
Decoding results during the final long-term memory retrieval phase. The figure illustrates the temporal evolution of average location decoding accuracy a. for the four experimental conditions and b. for the condition contrasts. Panel a depicts the contrast between the decoding accuracy and chance level (0.25). The green and blue shaded regions highlight the significant clusters: (i) 520-836 ms for the prioritization condition; (ii) 88-1100 ms for the testing condition; (iii) 1028-1200 ms for the absent in working memory condition. Panel b shows the three conditions contrasts. The prioritization vs. absent in working memory condition revealed a significant cluster between 472-640 ms and the testing vs. absent in working memory condition contrast an effect between 520-688 ms, as indicated by the grey shaded regions. The shaded area surrounding the decoding accuracy time series represents the standard error of the mean.

In the second step, we compared decoding accuracy between these conditions. The prioritization vs. absent in working memory condition comparison showed a significant cluster between 472-640 ms, while the testing vs. absent in working memory condition contrast showed a significant effect between 520-688 ms. No significant differences were found when comparing the prioritization vs. testing condition (Figure 4b). These results align with the parietal old-new pattern, indicating enhanced long-term memory representation for both the prioritization and testing condition compared to the absent in working memory condition.

## 3. Discussion

In two experiments, we investigated whether attentional prioritization (Experiments 1 and 2) and testing (Experiment 2) in working memory enhance long-term memory representations. A unique aspect of this study was examining the impact of these working memory processes on *existing* long-term memory representations that have been recently encoded but not yet consolidated. Additionally, in contrast to previous studies investigating comparable processes ^21–23,25,26^ we also explored these effects at a neural level. Behavioral and EEG results from Experiment 1 demonstrated a clear long-term memory retrieval benefit from attentional prioritization in working memory compared to the: (i) absence of information in working memory and (ii) brief presentation and maintenance in working memory (non-prioritization condition; also analogous to a repetition effect). Experiment 2 further revealed two important patterns of results.

First, results of Experiment 2 suggested that both the combination of attentional prioritization and testing (i.e., prioritization condition) and also testing in working memory (i.e., testing condition) led to long-term memory benefits relative to a control condition, which involved the absence of information in working memory. Importantly, in this context, the testing effect was defined as the non-selective maintenance of information in working memory to generate a response during the final match-no-match test. Moreover, in contrast to previous studies, we provide evidence for a boost in long-term memory retrieval both at the behavioral and at the neural level (ERP and decoding results). These findings add to previous literature suggesting that different processes in working memory (i.e., elaboration, attentional prioritization, testing) can lead to a differential built-up of information in long-term memory^9–15,17–26^. However, the present study offers a new perspective in this area, because it reveals that even recently encoded, unconsolidated long-term memories benefit from processing in working memory. Importantly, these benefits occur even when working memory processing is implicit. Participants were instructed to treat the working memory phase separately from the long-term memory task, and many were unaware that the objects appeared in the same locations during the encoding phase and subsequent working memory phase (cf., answers in the follow- up questionnaire). Thus, the observed enhancements in long-term memory retrieval are unlikely to be attributed to consistent strategic efforts by participants to reinforce previously encoded associations.

A key question in this context is understanding the mechanisms through which these two conditions enhance long-term memory retrieval compared to the absent in working memory condition. Several working memory processes likely contribute to this effect. First, in the prioritization condition, the selective retro-cue leads to the prioritization and selective maintenance of the cued item until the test. Consistent with previous research^14,36^, the presentation of the selective retro-cue enhances the object’s representation and its report, which in turn leads to the long-term memory retrieval benefit. In contrast, the testing condition did not involve attentional prioritization but instead relied on the non-selective maintenance of items until the working memory test.

Regarding the testing condition, we had two important observations. First, our results suggest that even though the working memory test did not require a location report, but rather a match no-match decision, this still benefitted long-term memory retrieval. This shows that long-term memory retrieval can also be boosted via testing in working memory, even though the working memory test and the final long-term memory test do not match. Second, results reported in the Supplementary materials (section 1, Figure S1) suggest that objects in the testing condition received a boost when presented as the central probe. Accordingly, items in the testing condition that were never presented as probes resulted in significantly lower accuracy compared to items that served as probes twice. Overall, this suggests that a non-specific working memory test can strengthen long-term memory representations, likely due in part to the additional boost items receive when serving as the central probe.

A second important finding of Experiment 2 is the dissociation between the behavioral and neural patterns. Behavioral evidence suggested a robust increase in accuracy when both attentional prioritization and testing were involved, compared to when only testing was involved. A similar pattern was observed in response times, although the evidence was weaker. At the neural level, a different pattern emerged. ERP results did not reveal a difference between the prioritization and testing condition. However, when these two conditions were contrasted with the absent in working memory condition (absence of information in working memory and mere repetition of information in working memory), the prioritization condition differed from both the absent in working memory condition, whereas the testing condition only differed from the absent in working memory condition. Finally, decoding retrieved locations from the EEG activity during the long-term memory task indicated no differences in the long-term memory representation between the prioritization and testing condition.

These findings from Experiment 2 indicate that while attentional prioritization provided a behavioral benefit over testing for long-term memory recall, this benefit was not mirrored at the neural level. In our view, this discrepancy might be due to the short-lasting nature of the neural benefits from attentional prioritization, which may not be detectable with EEG at the scalp level. This suggests that other brain imaging techniques, such as fMRI or intracranial EEG, may be needed to capture these subtle changes in neural activity.

Bringing together our findings, we argue that our results support the notion of attentional prioritization and testing in working memory jointly enhancing the retrieval of long- term memory content. This conclusion aligns with theoretical proposals from the working memory literature that suggest a tight link between attentional prioritization and testing. Accordingly, the prevailing view is that attentional prioritization in working memory always serves a specific purpose, such as preparing for a final test or performing an action^37–39^. Prioritizing and selecting information in working memory thus involves keeping the information in a heightened state, protected from interference until it serves its purpose^40^. Even unprioritized information can be restored to a prioritized state, if the participant is aware that this information is relevant for a subsequent test^41^. Consequently, information is only discarded from working memory when participants are confident that it will not be needed later, suggesting that attentional prioritization is driven by task-specific demands. Therefore, testing in working memory provides a functional purpose for attentional prioritization, indicating that these processes operate together to support effective memory retrieval and task performance.

At a broader level, our findings have significant implications for current theories of long-term and working memory. Traditional views suggest that long-term memory retrieval is mediated by working memory, which temporarily maintains retrieved long-term memory content^3,4,42^. Our results expand on these theories by indicating that working memory not only serves as a temporary buffer, but it is an instance, where active processes, such as attentional prioritization or testing can modify and strengthen the recently encoded long-term memory representations. This suggests that the relationship between the two memory systems is far more dynamic than previously thought, with working memory playing an active role in shaping long-term memory representations that were recently encoded and are not yet fully consolidated.

### Conclusions

To summarize, our study provides novel insights into the interplay between working- and long- term memory. Across two experiments, we collected data from 86 participants and demonstrated that attentional prioritization and testing within working memory *jointly* enhance the retrieval of existing long-term memory representations, with evidence at both behavioral and neural levels. Our findings extend existing theoretical views by suggesting that working memory does more than temporarily store retrieved long-term memory information; it can actively strengthen it. By demonstrating that recently encoded, not yet fully consolidated long-term memories benefit from working memory processes, our study highlights the critical role of working memory in shaping durable long-term memory traces.

## 4. Methods

### 4.1. Experiment 1

#### 4.1.1. Participants

Initially, data from 49 participants were collected, with six datasets excluded: four due to technical issues with EEG recording and two due to misunderstandings of the task. One of the latter participants consistently pressed the right button during the final retrieval phase, failing to report any actual location. The second participant was excluded based on the verbal feedback provided after the experiment, which revealed that the person misunderstood of the task. The final sample comprised 43 participants (25 females, 18 males) aged 18 to 31 years (*Mage* = 23.65 years, *SDage* = 2.85). We based our sample size on a previous investigation, in which we investigated processes unfolding during long-term memory retrieval^45^. All participants were right-handed, had normal or corrected-to-normal vision, and reported no history of neurological or psychiatric disorders. Compensation was provided either as a payment of 12 €/hour or as study participation credits for psychology students. Written consent was obtained prior to participation. The study was approved by the ethics committee of the Leibniz Research Centre for Working Environment and Human Factors (Dortmund, Germany) in accordance with the Declaration of Helsinki.

#### 4.1.2. Experimental procedure

Upon arrival, participants were given an overview of the study and written consent was obtained. Afterwards, they completed the German versions of a demographic survey and the Edinburgh Handedness Inventory^43^. Next, the EEG cap was prepared, and participants were guided into the dimly lit EEG laboratory. The experiment was run on a 22-inch CRT monitor (100 Hz; 1024×768 pixels) positioned at a viewing distance of ∼145 cm. The experimental task was programmed in Lazarus IDE (Free Pascal) and presented using the ViSaGe MKII Stimulus Generator (Cambridge Research Systems, Rochester, UK). The procedure began with a short training session, which was a short simulation of the main experiment.

The training consisted of 44 images (64 trials). This was followed by the main task, which consisted of a combination of a long-term and working memory task with three phases: an encoding phase, a working memory task, and a retrieval phase. Each phase was separated by a five-minutes break. At the conclusion of the experiment, participants completed a follow-up questionnaire, which asked about the strategies used by participants and encountered difficulties. The session typically lasted between 3 and 4.5 hours.

#### 4.1.3. Stimuli and materials

Stimuli for Experiment 1 were selected from a stimulus set comprising everyday objects^44^. Although the original pool included 260 objects, our selection was refined based on a prior image rating survey^45^, in which we selected a subset of 240 objects exhibiting optimal luminance, contrast, vividness, and recognizability. Thus, for each participant, a random subset of 180 images was selected for the task.

#### 4.1.4. Experimental paradigm

During the encoding phase, participants viewed an everyday object (size: 253×177px or 177×253px, corresponding to 4.8°×3.35° or 3.35°×4.8° visual angle) on a gray background (RGB 128-128-128) for 500 ms. The object appeared in one of four screen locations: top, bottom, left, or right. Participants were instructed to memorize the object’s location while keeping their gaze fixed on a central black dot (size: 0.2°×0.2°), positioned ∼3° from the object’s center (see Figure 1c). To aid memorization, they visualized the object in its location after it disappeared, still focusing on the central dot, and were instructed not to use verbal repetition or semantic information. At the end of each trial, participants confirmed memorization by clicking the right button on the response device, initiating the next trial. Each object was presented twice in the same location to facilitate learning, with all 180 objects shown once before any were repeated.

The second phase of the task involved a working memory retro-cue paradigm. Participants viewed a memory display for 300 ms, showing two objects from the encoding phase (size: 253×177px or 177×253px, corresponding to 4.8°×3.35° or 3.35°×4.8° visual angle) and two scrambled objects (size: 162×162px, corresponding to 3.07°×3.07° visual angle) around a fixation dot (size: 0.2°×0.2°). The center of each object was ∼3° from the fixation dot. The two real objects appeared in their original encoding locations. The scrambled objects were created using the GIMP software (https://www.gimp.org/) with the fx_taquin command from the G’MIC plugin (https://gmic.eu/), resized to 162x162px before scrambling. Participants were instructed to memorize the two real objects for a later report and ignore the scrambled objects, which only acted as sensory fillers.

After a 500 ms maintenance period, a retro-cue (size: 1°×0.5° or 0.5°×1°) indicated with 100% validity which item would be tested. The cue specified either the vertical or horizontal axis. After an 800 ms interstimulus interval, participants saw a central probe object and indicated whether it matched the cued object by pressing the top button for yes or the bottom button for no, with response mapping counterbalanced across subjects. The central probe matched the cued object 50% of the time (requiring a ‘yes’ response), it was a non-cued object 25% of the time and a new object in 25% of the time. Overall, participants were supposed to answer with a ‘no’ response in 50% of the trials.

Importantly, only 2/3 of the objects from the encoding phase were included in the working memory task: one-third (60 objects) were consistently cued (prioritized), another one-third (60 objects) appeared as non-cued / non-prioritized, and the remaining one-third appeared only in the encoding and final retrieval phases (see Figure 1d). In each trial, a prioritized and a non-prioritized object were randomly paired, ensuring one appeared on the horizontal axis and the other on the vertical axis. Each object, bound to its location and condition, appeared four times during this phase, resulting in 240 trials in total.

During the final retrieval phase, participants were shown all the objects from the encoding phase (size: 253×177px or 177×253px, corresponding to 4.8°×3.35° or 3.35°×4.8° visual angle). Their task was to report the location associated with each object during encoding. They used a response device with four buttons mapped to the possible locations: top, bottom, left, and right. Participants were instructed to keep their index finger on the middle of the device between trials and return to this position after responding. To increase the number of trials, each object’s location was reported twice, resulting in a total of 360 trials (180 objects, each presented twice).

#### 4.1.5. Data analyses

MATLAB® (R2021b) was utilized for conducting all behavioral, EEG, and statistical analyses. Initially, the dataset of each participant was segmented into three phases of the task: encoding phase, working memory task, and retrieval phase. Since we were interested in the consequences of differential processing in working memory, behavioral and EEG data from the retrieval phase was analyzed.

##### 4.1.5.1. Statistical analyses

We conducted our statistical analyses both using a frequentist and a Bayesian framework. The former analyses were conducted using the functions of the Statistics and Machine Learning Toolbox implemented in MATLAB® (R2021b). In case of the rm-ANOVA, Mauchly’s test for sphericity was applied. Whenever the sphericity assumption was violated, the Greenhouse– Geisser correction (represented by ε) was employed. Partial eta squared (ηp²) was calculated as the measure of effect size. For the t-tests applied, unless otherwise noted, two-tailed p-values were computed. Effect sizes were calculated using Cohen’s dav, following Lakens’ recommendations^46^. For the post-hoc tests requiring multiple comparison adjustments, false discovery rate ^47^ corrected *p*-values (denoted as *padj*) were provided .

The Bayesian statistical analyses were conducted in JASP (v.0.18.3)^48^. Within the Bayesian framework, Bayes factors denote the amount of evidence in favor of the alternative or the null hypothesis. BF10 represents evidence in favor of an effect or condition difference, while BF01 (1/BF10) denotes evidence for the lack of an effect or condition difference. Bayes factors lower than 3 have been interpreted as weak evidence, values between 3-20 as positive or substantial evidence, Bayes factor ranging between 20-150 as strong evidence, and finally values higher than 150 as very strong evidence^49^. For both the rm-ANOVA and the Bayesian t- tests, a multivariate Cauchly prior was adopted (as implemented in JASP). In case of the post- hoc t-tests, corrected posterior odds are reported, which are denoted by BF01 corr or BF10 corr.

##### 4.1.5.2. Behavioral analyses

In order to answer our main research question, accuracy and response times recorded during the retrieval phase were assessed. These two parameters were contrasted between the three experimental conditions: (i) prioritization condition, which includes objects that were cued and have undergone attentional prioritization in the working memory task; (ii) non-prioritization condition, which includes objects that were present in the working memory task but did not undergo attentional prioritization; (iii) absent in working memory condition, which includes objects that were absent in the working memory task. Statistical comparisons were conducted using a repeated measures analysis of variance (rm-ANOVA) with the within-subject factor condition.

##### 4.1.5.3. Preprocessing

The EEG data underwent preprocessing using the EEGLAB toolbox (version 14.1.2b; Delorme & Makeig, 2004) implemented in MATLAB®. The first step of the preprocessing pipeline was the application of a 0.1 Hz Hamming windowed sinc FIR high-pass filter (filter order: 33001, transition bandwidth: 0.1 Hz, cutoff frequency at −6 dB: 0.05 Hz) and a 30 Hz low-pass filter (filter order: 441, transition bandwidth: 7.5 Hz, cutoff frequency at −6 dB: 33.75 Hz) using the pop_eegfiltnew function. Following this, channels exhibiting substantial noise were identified and discarded through the automated channel rejection procedure in EEGLAB (for an overview, see Table 1) (pop_rejchan function). As a next step, data were re-referenced to the average. Notably, channels capturing anterior eye movements were excluded from rejection to optimize eye-movement-related component detection in the subsequent independent component analysis (ICA). Preparing for ICA, the data were downsampled to 200 Hz and subjected to a 1 Hz Hamming windowed sinc FIR high-pass filter (pop_eegfiltnew, filter order: 661, transition bandwidth: 1 Hz, cutoff frequency at −6 dB: 0.5 Hz). Subsequently, epochs spanning -1000 to 3000 ms relative to object onset were established, followed by baseline correction (-200 to 0 ms). Trials exhibiting extreme fluctuations were then rejected via EEGLAB’s automated trial rejection procedure (for an overview, see Table 1) (pop_autorej; threshold: 500 μV, maximum % of rejected trials: 5%). Next, ICA was conducted on the rank- reduced data (remaining number of channels minus one, determined via principal component analysis within the pop_runica function). Identification of artifact-containing independent components (ICs) was realized via the ICLabel plug-in (version 1.3; Pion-Tonachini et al., 2019), which categorizes ICs into brain, muscle, eye, heart, line noise, channel noise, and other noise. Following the criteria established by Wascher and colleagues (2022), ICs labeled with ≥ 30% probability as eye movement components or < 30% probability as brain components were discarded. Subsequently, the ICA weights were back-projected to the original 1000 Hz data, followed by band-pass filtering and re-referencing. Data were again epoched (-1000 to 3000 ms) and underwent baseline correction (-200 to 0 ms). ICs marked for rejection were removed, and trials with significant fluctuations were discarded using the same automated procedure mentioned above (for an overview, see Table 1) (threshold: 1000 μV, maximum % of rejected trials: 5%). Finally, missing channels were interpolated using the spherical spline method in EEGLAB (pop_interp).

##### 4.1.5.4. Event-related component analysis

In order to test our main hypothesis, we assessed the left parietal old-new ERP component, previously argued to capture long-term memory recollection ^28–32^. First, data was downsampled to 250 Hz. Next, we extracted the ERPs relative to the experimental conditions and data were averaged across the left parietal electrode cluster (P1, P3, P5)^34^. For the statistical comparison, a cluster-based permutation procedure was adopted. This procedure was applied three times for the (i) prioritization vs. non-prioritization condition; (ii) prioritization vs. absent in working memory condition; and (iii) non-prioritization vs. absent in working memory condition contrasts. For each analysis, the time window was restricted to 0-1200 ms after object onset. The typical left-parietal old-new effect usually occurs between 500-800 ms^32,34^; however, since we adopted cluster-based permutation statistics, the time window was extended.

The first step of the cluster-based permutation procedure was to contrast the time series of the two conditions using paired-sample t-tests. This resulted in a vector of 875 *p*-values, each corresponding to a comparison from the original dataset. Clusters with more than one significant *p*-value (*p* < 0.05) were identified based on this vector. In the next phase, a distribution of maximum cluster sizes was created by randomly shuffling the condition labels (e.g., prioritization and non-prioritization) within 1000 iterations. For each iteration, paired-sample t-tests were performed at all time points to compare the voltage changes between each condition. At each iteration, the largest cluster of significant *p*-values (*p* < 0.05) was recorded, which resulted in a distribution of maximum cluster sizes. Finally, we determined the 95th percentile of this distribution, and clusters exceeding this threshold were deemed significant.

### 4.2. Experiment 2

#### 4.2.1. Participants

Data were initially collected from 45 participants, with two participants subsequently excluded. One was removed for performing at chance level (25%) during the final retrieval phase, and the other due to having mean reaction times that were six standard deviations away from the group’s average. The final dataset consisted of 43 individuals. The participants (25 females, 18 males) ranged in age from 20 to 34 years (*Mage* = 24.97 years, *SDage* = 3.94). All other details from Experiment 1 are also applicable to Experiment 2.

#### 4.2.2. Experimental procedure

Experiment 2 included an experimental procedure identical to Experiment 1, with the only difference that the training consisted of more trials and images (52 trials and 15 images) and it was a simulation of the paradigm of Experiment 2.

#### 4.2.3. Stimuli and materials

Since Experiment 2 contained a new condition, which required significantly more objects (i.e., 300 in total), stimuli from the pool introduced by Brady and colleagues^53^ was used. In total, 300 objects were manually selected.

#### 4.2.4. Experimental paradigm

In Experiment 2, the encoding and retrieval phases were the same as in Experiment 1, but with 300 objects total, resulting in 600 trials for both phases. Due to using images from a different dataset, both the target and scrambled objects were sized at 200 px×200 px (corresponding to a visual angle of 3.8°×3.8°). A key difference from Experiment 1 was the introduction of a new condition in the working memory task: a neutral cue (size: 1°×1°) condition (referred to as testing condition) that provided no information about the to-be-tested item. Participants had to determine whether any object from the original memory display matched the probe object. Similar to Experiment 1, 50% of the probes were objects from the display, and 50% were entirely new objects, never encountered in the task. In Experiment 2, objects were divided into four conditions (see Figure 1d): (i) one-fifth were in the cued (prioritization) condition; (ii) one-fifth were in the non-cued (non-prioritization) condition; (iii) two-fifths were in the testing condition; and (iv) one-fifth were in the absent in working memory condition, not appearing in the working memory task.

#### 4.2.5. Data analyses

All aspects reported in case of Experiment 1 apply also for Experiment 2.

##### 4.2.5.1. Behavioral analyses

In Experiment 2, we aimed to compare final long-term memory retrieval performance— measured by accuracy and response times—across four experimental conditions using rm-ANOVA. These conditions were: (i) prioritization condition, where objects were cued and underwent attentional selection and testing in the working memory task; (ii) non-prioritization condition, where objects were present in the working memory task but did not undergo attentional selection or testing; (iii) absent in working memory condition, where objects were absent from the working memory task; and (iv) testing condition, where objects were cued and tested but did not undergo attentional selection (Figure 1d). Since Experiment 2 included more objects and thus participants were required to learn more associations, we conducted a control analysis to ensure that performance in Experiment 1 and 2 was comparable (see Supplementary materials, section 3, Table S3).

##### 4.2.5.2. Preprocessing

The preprocessing pipeline adopted for the ERP analysis of Experiment 2 was identical to the pipeline adopted in case of Experiment 1. Nevertheless, in case of the decoding analysis, we introduced some changes. First, except for a 1 Hz low-pass filter used to prepare data for the ICA, filtering was entirely omitted, as prior research suggests that filtering might artificially boost decoding results^54^. Furthermore, because we wanted to maximize the number of trials used for decoding, we opted to exclude the trial rejection procedure from this pipeline. As a result, we had 120 trials per each condition (for the prioritization, non-prioritization, and control condition), and 240 trials for the testing condition. With both filtering and trial rejection omitted, we implemented channel rejection across the full range of electrodes.

##### 4.2.5.3. Event-related potential analysis

We applied the same analysis described in case of Experiment 1. Since Experiment 2 includes four conditions, six different statistical comparisons were conducted, which enabled the contrasting all conditions with all the other conditions.

##### 4.2.5.4. Decoding analyses

Our analysis was centered on decoding the location retrieved during the final long-term retrieval phase. First, we performed decoding within each condition and then compared the decoding accuracy between conditions – except the non-prioritization condition, which did not show above-chance decoding accuracy. Specifically, the classifier was trained to distinguish between one vs. all the other retrieved locations (e.g., top location vs. all the other locations) within each condition. Thus, the classification routine was run for each condition separately.

Following the methodology of Bae and Luck^55^, we used a combination of support vector machines and error-correcting output codes, implemented with the fitcecoc() MATLAB® function, while testing and predictions were performed using the predict() function.

The decoding procedure’s input consisted of power values from time-frequency decomposition. We included twenty-six frequency bands between 4-30 Hz and 64 electrodes as features. Classification was conducted separately for each participant and timepoint, focusing on EEG oscillatory activity between -200 to 1300 ms. At each timepoint, trials for each location were randomly divided into three groups, referred to as blocks. An equal number of trials for each location were assigned to each block, and any extra trials were removed if the total number could not be divided by three. Within each block, data were averaged for each location label, resulting in three averages per location. We used trial averages instead of single trial data, as previous investigations argued that averaging reduces noise^55^. This also ensured that results were not due to decoding object identity. Model training was performed on 2/3 of these averages, with testing on the remaining 1/3 (i.e., 3-fold cross-validation). Each average served once as the test dataset. Once all averages served as a test dataset, trials were reshuffled and re-assigned to the blocks for averaging and this procedure was repeated for 10 iterations. Predicted labels for all timepoints and iterations were compared to true labels to calculate decoding accuracy. This was obtained and subsequently averaged across iterations. A five-point moving time window smoothed these time series, and decoding accuracies were averaged across participants. Since four locations were contrasted, the chance level was 25%.

First, we compared the decoding accuracy within each condition to chance level (0.25) and next we compared it between conditions, using the cluster-corrected sign-permutation test adopted by Wolff and colleagues ^56^. This method generates a null distribution by flipping value signs with a 50% probability over 100 000 iterations (function: cluster_test_helper). Significant clusters in the actual data were identified using this null distribution (function: cluster_test), with significance thresholds set at *p* < .05. Based on the outcome of Experiment 1, we hypothesized that the decoding accuracy should be greater than chance level and that the decoding accuracy in the prioritization and testing condition should be higher than that in the absent in working memory condition, one-tailed, cluster-corrected sign permutation tests were used. Comparable to the ERP analysis, the statistical tests were limited to the 0-1200 ms window.

## Acknowledgments

The authors would like to thank Tobias Blanke for programming the experiment and providing technical support; Nathalie Hutterloh for assistance with data collection, participant recruitment, and lab organization; and all the students who contributed to the data collection process: Amina Aljandali, Madlen Bernards, Niklas Burczyk, Oliver Butzke, Alper Capakli, Quang Dang, Patrick Frenken, Franziska Göpp, Lorenza Schlumbom, Pia-Lotta Schmidt, Vivien Szczepanski, and Stefan Weber.

## Author contribution

M.S.: data collection and data analysis. M.S. and D.S.: conceptualisation. M.S., L.I.K., D.S.: writing the original draft. D.S.: project supervision and acquiring the necessary funding.

## Open practices and data availability statement

The experiment was not preregistered. The raw data and all the scripts used for the reported analyses will be publicly available after publication on Open Science Framework (OSF): https://osf.io/ne9yt/

