## Supplementary materials for "Working memory processing boosts the neural representation of long-term memories"

#### 1. Long-term memory accuracy changes as a function of probing instances in the testing condition

To further explore the impact of probing during the working memory phase on subsequent long-term memory retrieval performance, we conducted an additional set of behavioral analyses. These analyses focused on the number of probing instances during the working memory task, defined as the frequency with which a specific object was used as a probe (i.e., appeared in the center of the screen during the final test of the working memory task) within a given condition. This analysis was feasible because, during the working memory phase, each object appeared in four different trials. Given that the neutral cue condition included a total of 240 objects, we were able to distinguish between trials with objects assigned to the neutral cue condition that were probed zero times, once, or twice.

For the accuracy comparison, we first checked whether the sphericity assumption was violated. Because this was not the case, we conducted a rm-ANOVA (factors: presented as probe zero times, once, and twice), which revealed a main effect of probing, $F(2, 84) = 6.48$, $p = .002$, $\eta_p^2 = 0.13$, $BF_{10} = 12.95$, with the Bayesian analysis suggesting substantial evidence in favor of the alternative hypothesis. Our descriptive analysis (see Table S1) suggests that participants were the most accurate in trials, in which the object was twice the probe, followed by trials with the object was once the probe, and finally in trials, in which the object was zero times the probe. The post-hoc analyses revealed strong evidence in favor of the difference between the zero vs. twice probed condition, $t(42) = -3.85$, $p_{adj} = .001$, $d_{av} = 0.33$, 95% CI [-9.10, -2.84], $BF_{10\ corr} = 40.34$. The lack of accuracy difference between the zero vs. once probed was supported by substantial evidence, $t(42) = -1.54$, $p_{adj} = .12$, $d_{av} = 0.14$, 95% CI [-6.12, 0.81], $BF_{01\ corr} = 0.29$. Finally, the lack of accuracy difference between the once vs. twice probed was supported by weak evidence, $t(42) = -1.93$, $p_{adj} = .08$, $d_{av} = 0.18$, 95% CI [-6.77, 0.13], $BF_{01\ corr} = 0.53$ (see Figure S1a).

The same rm-ANOVA was also conducted for response times. No main effect of probing was found, $F\,(2, 84) = 2.63$, $p = .07$, $\eta_p^2 = 0.05$, $BF_{01} = 0.63$ and the Bayes factor indicated only weak evidence in favor of the null hypothesis (see Figure S1b).

**Table S1**

*Descriptive statistics*

**Mean and standard deviation**

| Probing number | N | Mean accuracy | SD accuracy | Mean response times | SD response times |
|---|---|---|---|---|---|
| Zero times | 43 | 68.70% | 17.34% | 1198.18 ms | 304.45 ms |
| Once | 43 | 71.36% | 18.44% | 1186.83 ms | 383.72 ms |
| Twice | 43 | 74.68% | 18.22% | 1123.57 ms | 296.96 ms |

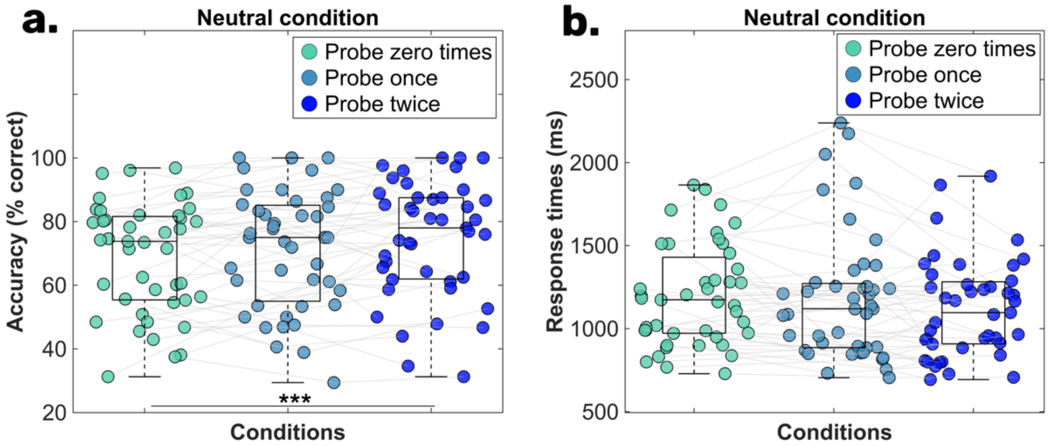


**Figure S1.** The scatterplots of panel a and b illustrate the mean accuracy and mean reaction time for objects that were a probe zero times, once, and twice. The central mark in each boxplot represents the median, while the bottom and top edges denote the 25th and 75th percentiles, respectively. The whiskers do not extend to averages, which are treated as outliers. Grey lines connect data points from the same participant. Statistical significance is indicated with the following symbols: * p < 0.05; ** p < 0.01, *** p < 0.001.

### 2. Higher accuracy in the prioritization condition when the number of probing instances constant

As a control analysis, we examined whether the observed accuracy differences between the prioritization and testing conditions held when the number of probing instances was kept constant. We compared accuracy between these conditions when objects were probed once and when they were probed twice. For the probed-once condition, our analysis revealed substantial evidence for a difference between the prioritization and testing conditions, $t(42) = 3.23$, $p = .002$, $d_{av} = 0.34$, 95% CI [2.29, 9.93], $BF_{10} = 13.70$. Similarly, when objects were probed twice, the accuracy difference was supported by strong evidence, $t(42) = 4.03$, $p < .001$, $d_{av} = 0.30$, 95% CI [2.61, 7.85], $BF_{10} = 113.83$ (see descriptive statistics in Table S2).

**Table S2**

*Descriptive statistics*

| | | | **Mean and standard deviation** | |
|---|---|---|---|---|
| **Probing number** | **Condition** | **N** | **Mean accuracy** | **SD accuracy** |
| Once | Prioritization | 43 | 77.47% | 16.93% |
| | Testing | 43 | 71.36% | 18.44% |
| Twice | Prioritization | 43 | 79.91% | 16.28% |
| | Testing | 43 | 74.68% | 18.22% |

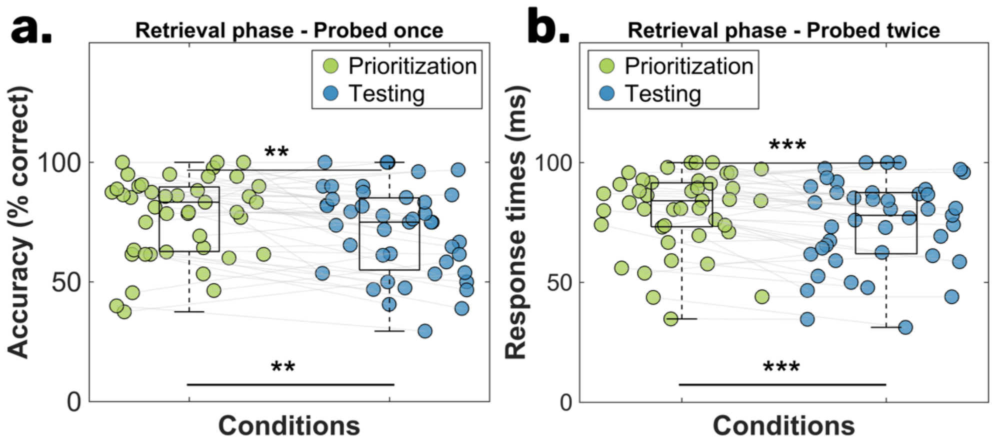


**Figure S2.** Panel a illustrates the mean accuracy in the prioritization and testing conditions for objects that served as a probe once during the working memory task, while panel b shows the accuracy difference for objects that served as a probe twice. The central mark in each boxplot represents the median, with the bottom and top edges indicating the 25th and 75th percentiles, respectively. The whiskers do not extend to averages, which are treated as outliers. Grey lines connect data points from the same participant. Statistical significance is indicated with the following symbols: * $p < 0.05$; ** $p < 0.01$; *** $p < 0.001$.

### 3. Comparable behavioral performance between Experiment 1 and 2

Given the higher number of objects to be learned in Experiment 2, we conducted an additional control analysis to ensure that behavioral performance in the final retrieval phase was comparable between experiments. To do this, we calculated the average accuracy and response times, then contrasted these measures using paired-sample t-tests (see Table S2). The results indicated no significant differences in accuracy, $t(42) = 0.60$, $p = .54$, $d_{av} = 0.12$, 95% CI [-0.04, 0.08], $BF_{01\ corr} = 0.19$ or response times, $t(42) = -0.47$, $p = .63$, $d_{av} = 0.10$, 95% CI [-173.70, 107.72], $BF_{01\ corr} = 0.18$ between the two experiments. Bayesian analysis further provided substantial evidence supporting the lack of performance differences.

**Table S3**

*Descriptive statistics – condition-independent behavioral performance of Experiment 1 and 2*

**Mean and standard deviation**

| Experiment | N | Mean accuracy | SD accuracy | Mean response times | SD response times |
|---|---|---|---|---|---|
| Experiment 1 | 43 | 72.90% | 14.89% | 1144.61 ms | 321.12 ms |
| Experiment 2 | 43 | 70.99% | 15.92% | 1177.60 ms | 300.90 ms |
